# Novel Muscle-Tropic AAV Capsids with Dramatically Enhanced Transduction and Safety Profiles in Non-Human Primates

**DOI:** 10.64898/2026.05.22.727076

**Authors:** Youguang Luo, Linlin Zhang, Zhenhua Wang, Hao Li, Rui He, Xiaofei Lv, Xianggang Xu, Shengwen Wang, Zhen Sun, Mengmeng Yu, Qun Zhang, Peilu Zhao, Long Wang, Bo Sun, Daoyuan Li, Zhenming An

## Abstract

Adeno-associated virus (AAV) gene therapy holds immense promise for treating muscular dystrophies, yet its efficacy and safety are constrained by the suboptimal tissue tropism of natural serotypes. Here, we employed the REACH platform, which combines rational design and directed evolution, to engineer muscle targeting vectors. Systemic administration in non-human primates (NHPs) revealed that lead candidate M1 mediates a >10-fold increase in skeletal muscle transduction compared to the AAV9 and 2-3 fold higher than MyoAAV, while concurrently achieving a remarkable 183-fold reduction in liver distribution. Furthermore, M1 exhibited significant de-targeting from key off-target tissues, including dorsal root ganglia (11 fold), lung (27 fold), spleen (2 fold), and kidney (2 fold). These findings demonstrate that the REACH platform can generate AAV capsids with simultaneously enhanced muscle tropism and favorable safety profiles, addressing a critical bottleneck in muscle-directed gene therapy.

## INTRODUCTION

Recombinant adeno-associated virus (AAV) is the leading vector for in vivo gene therapy.^1,2^ For systemic treatment of widespread muscle diseases such as Duchenne muscular dystrophy (DMD), achieving high and specific transduction of skeletal and cardiac muscle is paramount.^3-5^ While serotypes like AAV9 exhibit some muscle tropism, their significant sequestration by the liver and transduction of other off-target tissues necessitate high, potentially toxic doses and raise safety concerns.^6,7^

Engineering novel AAV capsids with improved targeting and reduced immunogenicity is therefore a major focus of the field.^3,4,6^ Current engineering strategies, including directed evolution and structure-guided design, have yielded progress but often face trade-offs between efficacy, specificity, and manufacturability.^8,9^ We developed the REACH platform to systematically overcome these limitations. This report details the application of REACH to generate a new class of muscle-tropic AAV vectors. We present comprehensive NHP data showing that the lead candidate, M1, not only surpasses AAV9 in muscle transduction efficiency but also establishes a new benchmark for systemic safety through profound liver de-targeting.

## RESULTS

### REACH Platform for AAV Capsid Engineering

The REACH (Rational and Evolutional AAV Customization/Creation Hub) platform is a proprietary, integrated pipeline for developing next-generation engineered AAV delivery vectors. It combines rational design with directed evolution through the construction of capsid libraries with large capacity and high diversity. This approach maximizes the potential for discovering variants with novel and superior biological properties. The platform workflow encompasses large-capacity library construction, iterative in vitro screening under selective pressures for efficacy and safety, followed by stringent in vivo screening in non-human primate (NHP) models. Final lead candidates are then advanced through confirmatory studies in NHPs, and further advanced toward clinical translation (Figure 1).

**Figure 1.**
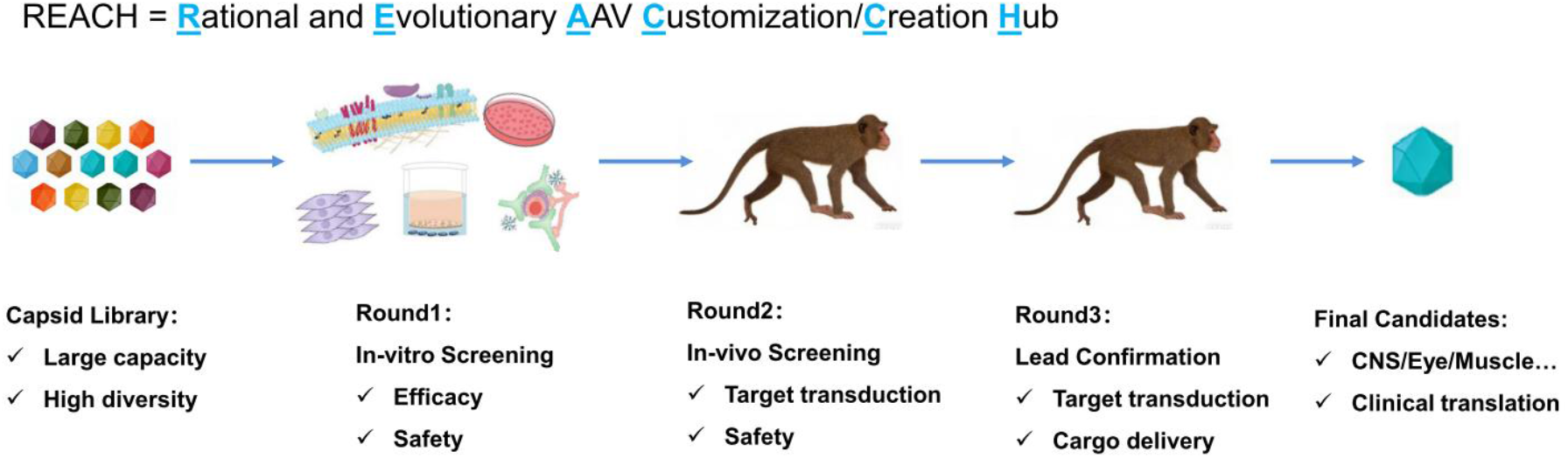
Schematic diagram of REACH platform for AAV engineering. REACH platform combined rational design with directed evolution. In general, a capsid library with large capacity and high diversity was constructed. And then after in-vitro screening, in-vivo screening and lead confirmation, final candidates would be found and subsequently advanced to clinical translation.

### Identification of Muscle-Tropic Candidates via In Vitro Screening

This capsid library was subjected to several in vitro screening designed to probe different mechanisms relevant to efficacy and safety. Here we show the representative efficacy screening with RGD-related receptor binding mechanism. In this part, the variants were screened with validated αVβ6 stable cells for transduction efficiency (Figure 2A and 2B). The promising variants with better performance than AAV9 were selected for subsequently screening (Figure 2C and 2D).

**Figure 2.**
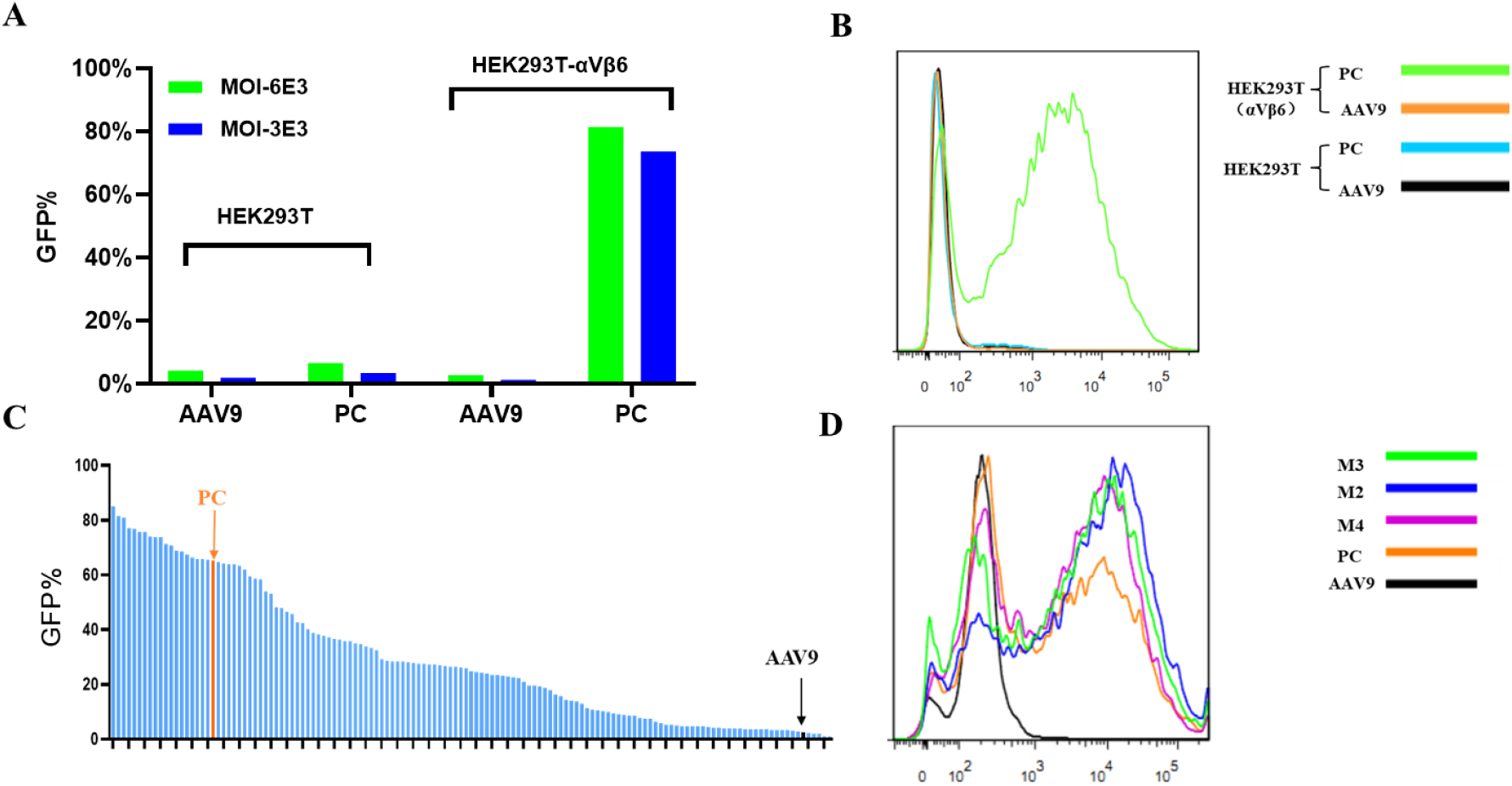
Representative in vitro efficacy screening. (A) αVβ6 stable cell line validation. Stably αVβ6-overexpressing HEK293T cells were transduced with AAV9 and PC at multiplicities of infection (MOIs) of 6×10^3^and 3×10^3^, and the percentage of GFP-positive cells was analyzed with flow cytometry. (B) Flow cytometry profiles for HEK293T and αVβ6 stable cell line (MOI: 6×10^3^). (C) Flow cytometry analysis of the percentage of GFP-positive αVβ6 cells. AAV9 and engineered variants expressing EGFP were used to infect HEK293T-αVβ6 cells for 48 hours, and then cell samples were collected and analyzed with flow cytometry. (D) Representative flow cytometry data of lead variants. PC indicates positive control, MyoAAV.

### Lead Capsids Show Enhanced Muscle Transduction in NHPs

These selected candidates were packaged with a barcoded genome, pooled, and intravenously administered to NHPs. Tissues of interest, including skeletal muscle, heart, spleen, kidney, lung, liver, and DRG, were collected and analyzed with next-generation sequencing (NGS, Figure 3A).

**Figure 3.**
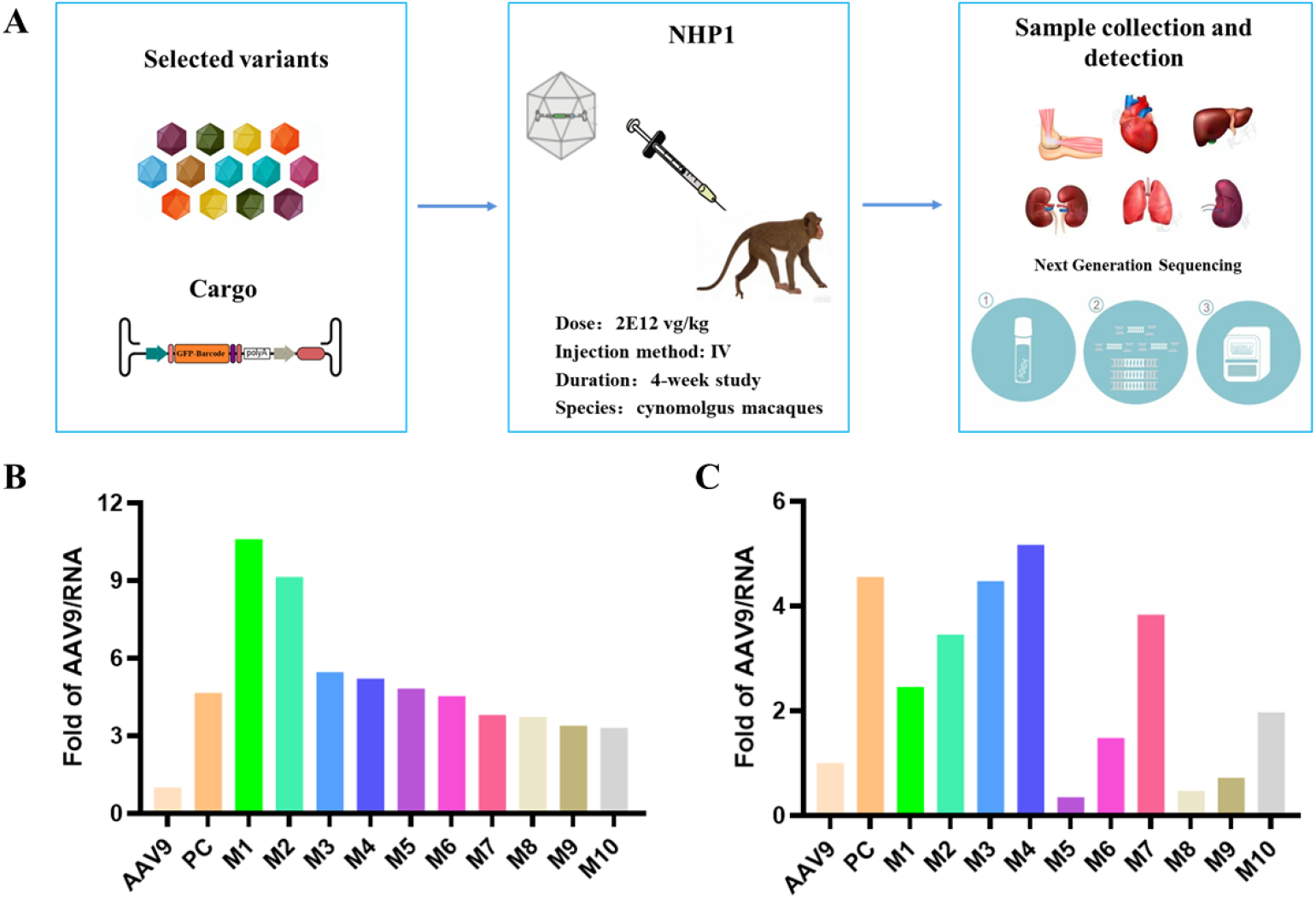
Engineered capsids exhibited enhanced transduction efficiency in skeletal muscle in NHPs. (A) Schematic diagram of in vivo screening in NHPs. Selected variants were injected intravenously to NHPs, and then samples of interest were collected and analyzed with NGS. (B) Fold change of AAV distribution in skeletal muscle compared to AAV9 in RNA level. (C) Fold change of AAV distribution in heart compared to AAV9 in RNA level. PC indicates positive control, MyoAAV.

The NGS results demonstrated that these candidate capsids mediated a 3-10 fold increase in skeletal muscle transduction compared to AAV9 and also 2-3 fold higher than that of positive control (MyoAAV),^5^ with different levels of heart transduction (Figure 3B and 3C). This establishes a clear efficacy advantage for muscle-directed gene delivery.

### Candidate Capsids Exhibit Superior Systemic Safety Profiles in NHPs

A critical finding was the dramatically improved safety profile of the REACH-engineered capsids. All lead candidates showed reduced distribution in non-target tissues compared to AAV9 (Figure 4 and Table 1).

**Table 1.**
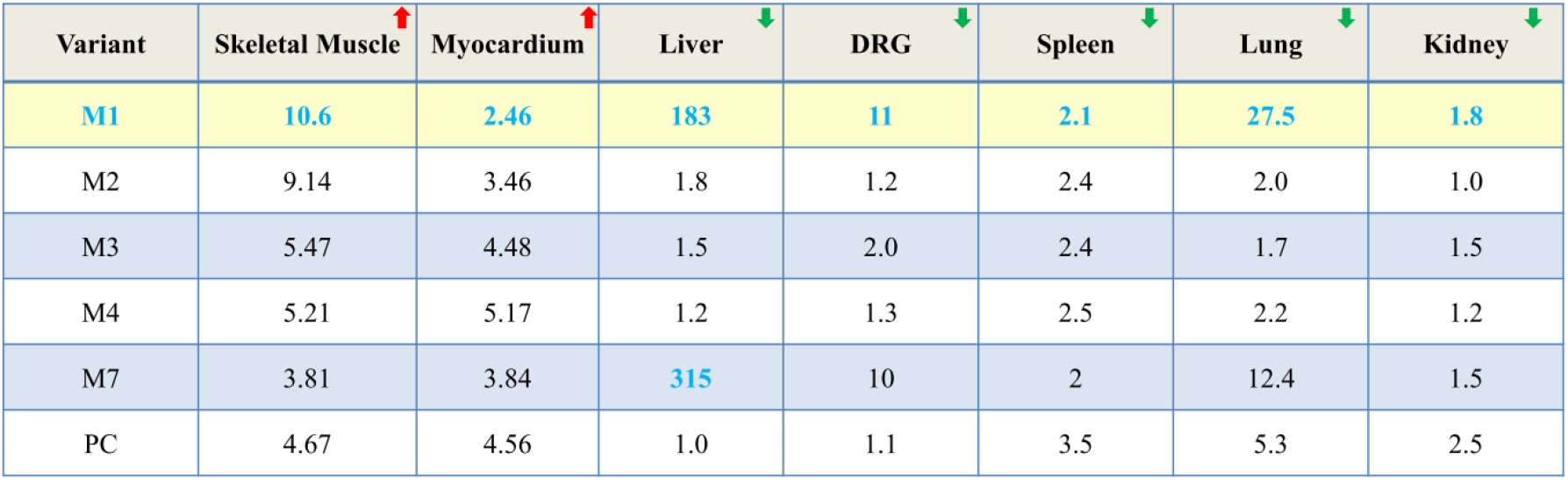
Safety profile of muscle-tropic delivery candidates.

| Variant | Skeletal Muscle ↑ | Myocardium ↑ | Liver ↓ | DRG ↓ | Spleen ↓ | Lung ↓ | Kidney ↓ |
| --- | --- | --- | --- | --- | --- | --- | --- |
| M1 | 10.6 | 2.46 | 183 | 11 | 2.1 | 27.5 | 1.8 |
| M2 | 9.14 | 3.46 | 1.8 | 1.2 | 2.4 | 2.0 | 1.0 |
| M3 | 5.47 | 4.48 | 1.5 | 2.0 | 2.4 | 1.7 | 1.5 |
| M4 | 5.21 | 5.17 | 1.2 | 1.3 | 2.5 | 2.2 | 1.2 |
| M7 | 3.81 | 3.84 | 315 | 10 | 2 | 12.4 | 1.5 |
| PC | 4.67 | 4.56 | 1.0 | 1.1 | 3.5 | 5.3 | 2.5 |

Table 1. Safety profile of muscle-tropic delivery candidates
| Variant | Skeletal Muscle | Myocardium | Liver | DRG | Spleen | Lung | Kidney |
| --- | --- | --- | --- | --- | --- | --- | --- |
| M1 | 10.6 | 2.46 | 183 | 11 | 2.1 | 27.5 | 1.8 |
| M2 | 9.14 | 3.46 | 1.8 | 1.2 | 2.4 | 2.0 | 1.0 |
| M3 | 5.47 | 4.48 | 1.5 | 2.0 | 2.4 | 1.7 | 1.5 |
| M4 | 5.21 | 5.17 | 1.2 | 1.3 | 2.5 | 2.2 | 1.2 |
| M7 | 3.81 | 3.84 | 315 | 10 | 2 | 12.4 | 1.5 |
| PC | 4.67 | 4.56 | 1.0 | 1.1 | 3.5 | 5.3 | 2.5 |

**Figure 4.**
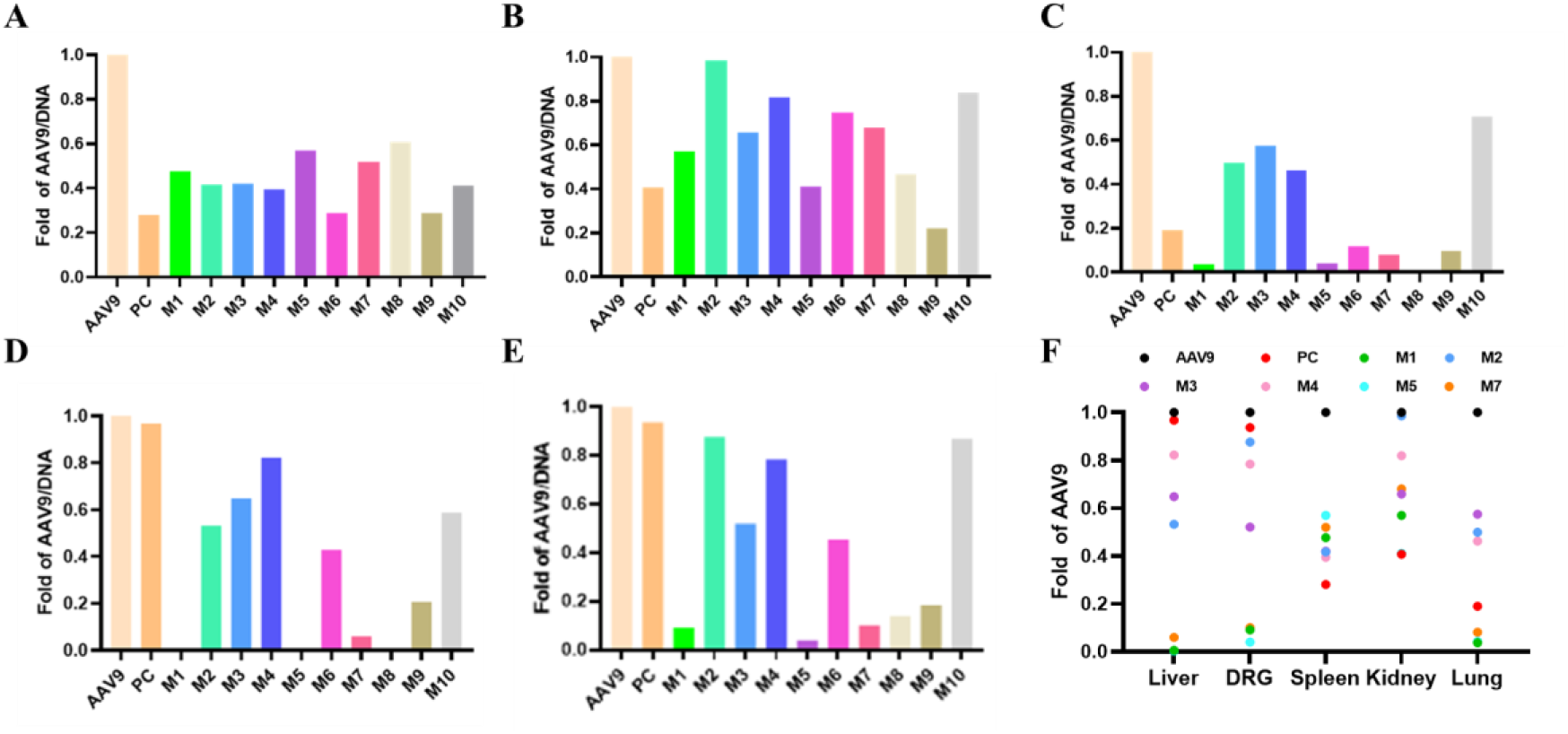
Engineered candidates show a favorable safety profile. (A) Fold change of AAV distribution in spleen. (B) Fold change of AAV distribution in kidney. (C) Fold change of AAV distribution in lung. (D) Fold change of AAV distribution in liver. (E) Fold change of AAV distribution in DRG. (F) Summary of fold changes of AAV relative to AAV9 in liver, DRG, spleen, kidney and lung. NHPs were intravenously injected with indicated AAV samples. Samples of interest were collected and analyzed with NGS, and the fold change of AAV variant relative to AAV9 was calculated. PC indicates positive control, MyoAAV.

Candidate M1 emerged as the standout, demonstrating a near-undetectable level in the liver, representing a 183-fold reduction versus AAV9 (Figure 4D). Furthermore, M1 showed significant de-targeting from other sensitive tissues: an 11 fold reduction in dorsal root ganglia (DRG), a 27 fold reduction in lung, and approximately 2 fold reductions in spleen and kidney distribution (Figure 4A-C, 4E and 4F). This comprehensive de-targeting profile suggests a substantially lowered risk of vector-related toxicities.

## DISCUSSION

This study introduces the REACH platform and its successful application in developing a novel muscle-tropic AAV vector, M1. The data robustly demonstrate that engineering can break the traditional efficacy-safety trade-off. M1’s >10-fold enhancement versus AAV9, and also 2-3 fold higher than the benchmark MyoAAV, in skeletal muscle delivery could directly translate to lower therapeutic doses, reducing manufacturing burdens and cost (Table 1). More importantly, its unprecedented liver de-targeting (183-fold reduction) directly addresses one of the most significant dose-limiting toxicities in systemic AAV therapy-hepatotoxicity and associated immune responses.^10-12^

The broad de-targeting profile, particularly from DRG and lung, is equally promising. DRG transduction has been linked to neurotoxicity in some preclinical studies, while lung off-targeting is undesirable.^13,14^ M1’s significant reduction in these tissues suggests a cleaner and potentially safer vector. The mechanistic basis for this simultaneous enhancement in muscle tropism and reduction in off-targeting likely involves altered interactions with cell surface receptors and/or glycan moieties, a subject of ongoing investigation.^9,15,16^

These candidates, particularly M1, are ideally suited for treating monogenic muscle disorders. For DMD, the high skeletal muscle transduction could improve the distribution and efficacy of micro-dystrophin genes. The retained heart tropism is beneficial for addressing cardiomyopathy. Furthermore, the favorable profile makes these vectors compelling delivery tools for muscle-targeted gene editing (e.g., CRISPR-Cas) or RNA-targeting platforms, where precise, efficient, and safe delivery is critical. And more clinical translation related studies need to be performed and identified, such as more detailed efficacy characterization, putative receptor identification, human translation analysis, comprehensive safety profile study, manufacturability evaluation.^17-20^

In conclusion, the REACH platform has generated AAV capsids that represent a significant advance over current benchmark MyoAAV.^5^ Candidate M1, with its potent muscle-specific transduction and exceptional safety profile in NHPs, is a prime candidate for clinical development. And the other candidate capsids with different skeletal and myocardial muscle transduction rate, could also be applied to different indications based on its delivery profile.^21,22^ This work underscores the power of integrated engineering platforms to create tailored gene therapy vectors that overcome the central delivery challenges in the field.

## MATERIALS AND METHODS

### AAV Vector Production and Purification for in vitro Validation Studies

All AAV vectors were produced using the standard triple-plasmid transfection method in suspension HEK293 cells (Thermo Fisher Scientific, VPCs2.0, A52021).

- **Cell Culture and Transfection:** HEK293 cells were maintained in CD05 medium (OPM Biosciences, P688293) and passaged when density reached ≥3×10^6^ cells/mL. For transfection, cells were seeded at 2×10^6^ cells/mL. A mixture of three plasmids—the adenoviral helper plasmid, the AAV rep/cap plasmid (wild-type or variant), and the transgene plasmid encoding enhanced green fluorescent protein (EGFP)—was combined at a 1:1:1 molar ratio. The DNA was mixed with polyethylenimine (PEI) transfection reagent (Yeasen, 40816ES03) at a PEI:DNA=2:1 ratio (v/w) and added to the cells.
- **Harvest and Lysis:** Three days post-transfection, cells were lysed by adding a lysis buffer containing Tris, MgCl_2_, and Tween-20, along with Benzonase (50 U/mL, Novoprotein, GMP-1707) to digest unpackaged nucleic acids. The lysate was incubated at 37°C for 3 hours and then clarified by centrifugation.
- **Purification:** The clarified harvest was filtered and subjected to affinity chromatography using an AAVX resin. The eluted virus was further purified by iodixanol (Shanghai Yuanpei, R714JV) density gradient ultracentrifugation. The final viral band was collected, buffer-exchanged, and concentrated using tangential flow filtration (TFF) or ultrafiltration spin column.

### In Vitro Transduction Assays

The engineered capsids were screened with several in vitro screening assays with different mechanisms representing efficacy and safety. Here is the brief introduction about RGD-based cell transduction assay.

- **Construction of the HEK293T-αVβ6 Cell Line:** Plasmids encoding the αV and β6 genes, along with a transposase plasmid, were co-transfected into HEK293T cells. After 24 hours, puromycin and hygromycin were added for dual selection to establish a stable cell line co-expressing the αVβ6 receptor. To confirm successful construction, both the generated HEK293T-αVβ6 and the parental HEK293T cells were infected with AAV9 or benchmark MyoAAV at various MOI. Successful transduction was assessed by measuring the GFP-positive rate via flow cytometry.
- **HEK293T-αVβ6 Cell Transduction:** The 293T-αVβ6 cells were seeded into 24-well plates at 1×10^5^ cells/well in DMEM complete medium (Gbico). Cells with 60-80% confluence were infected with AAV vectors at various MOIs in serum-free medium for 4 hours, followed by addition of complete medium to reach 10% FBS. After 48 hours, transduction efficiency was quantified by flow cytometer (BD Biosciences) to determine the percentage of GFP-positive cells and fluorescence intensity.

### Large-Scale Production and Evaluation in Non-Human Primates (NHPs)

- **Samples Production for NHP Study:** For the NHP study, each selected variant was individually packaged with a unique DNA barcode within the EGFP expression cassette. After titer determination, equal genome copies of each variant were pooled. The pooled library was purified with TFF, AAVX affinity chromatography, iodixanol density gradient centrifugation, and buffer exchange. The final product was assessed for titer, purity (by SDS-PAGE), capsid full/empty ratio, homogeneity (by DLS), endotoxin level, and sterility.
- **NHP Experimental Design:** The study was conducted in cynomolgus macaques (Macaca fascicularis). Prior to dosing, animals were screened for pre-existing neutralizing antibodies (NAbs) against AAV9, and qualified macaques with lower NAb titers were selected.
- **Dosing and Tissue Processing:** Each animal received a single intravenous infusion of the pooled AAV library at a dose of 2×10 ^12^ vg/kg. 28 days post-dosing, animals were perfused and necropsied. Tissues, including skeletal muscle, heart, liver, spleen, lung, kidney and DRG, were collected, snap-frozen, and stored at -80°C.
- **Biodistribution Analysis via NGS:** DNA and/or RNA were/was extracted from all samples as indicated. The region containing the unique barcode was amplified by PCR from each sample, and the resulting amplicons were subjected to NGS. The relative abundance of each barcode (corresponding to each AAV variant) in every tissue was calculated from the NGS read counts to determine the biodistribution profile.

### Statistical analysis

Unpaired two-tailed t tests were performed in GraphPad Prism and are reported in figures or figure legends. A p value <0.05 was considered significant. * indicates p<0.05, ** indicates p<0.01, and *** indicates P<0.001.

## DATA AVAILABILITY

The data that support the findings of this study are available from the corresponding author upon reasonable request.

## ACKNOWLEDGMENTS

This study was funded by Qilu Pharmaceutical Co., LTD. The authors thank the Institute of Biopharmaceuticals team, pre-clinical team, et al. for their contribution in designing/performing the experiments and data analysis.

## AUTHOR CONTRIBUTIONS

Conceptualization: YL, LZ, ZA; Methodology: LZ, ZW, HL, RH; Experimentation: LZ, ZW, XX, ZS, HL, RH, MY, SW, XL, QZ, PZ, LW; Resources: ZA, YL, BS, DL; Formal

Analysis: LZ, ZW, HL, RH, ZS, XX; Investigation: LZ, ZW, HL; Writing-Original Draft: YL, ZW, HL, XX, ZS; Writing-Review & Editing: YL, LZ, ZA.

